# Alcohol use disorder is associated with altered frontomedial phase-amplitude coupling strength during resting state

**DOI:** 10.1101/2025.05.13.653768

**Authors:** C.D. Richard, B. Porjesz, J. L. Meyers, A. Bingly, D.B. Chorlian, C. Kamarajan, G. Pandey, A. Anokhin, S. Brislin, W. Kuang, A. K. Pandey, S. Kinreich

## Abstract

Considerable evidence from functional neuroimaging and EEG coherence studies indicates that individuals afflicted with alcohol use disorder (AUD) manifest aberrant patterns of connectivity, particularly in frontal brain regions. Phase-amplitude coupling (PAC) is another form of functional connectivity, reflecting the association between the phase at one frequency and amplitude changes at a higher frequency. Significant PAC differences have been reported for other substance use disorders, but it has not yet been investigated in AUD. We compared frontomedial PAC strength during resting state, eyes closed, in adult participants with severe AUD and age-matched unaffected controls from the Collaborative Study on the Genetics of Alcoholism (COGA). Comodulograms of PAC estimates between phase frequencies (0.1-13 Hz) and amplitude frequencies (4-50 Hz) were calculated for all participants. PAC differences between AUD and unaffected groups were assessed at each phase-amplitude frequency pair in comodulograms to identify clusters of significant test results, reporting only those clusters satisfying all validation and significance testing steps. Severe AUD was associated with clusters of significantly greater PAC in alpha-gamma domains of both men and women. Candidate clusters were found in theta-gamma domains of both sexes, but were only significant greater in men with AUD. Significant PAC clusters were found in the delta-gamma domain of both sexes, though women with AUD showed significant decreases in contrast to greater PAC found in men with AUD. The significant PAC clusters identified in this exploratory study could provide new insights into the dysregulation of brain connectivity underlying AUD.

## INTRODUCTION

Alcohol use disorder (AUD) is characterized by compulsive tendencies for alcohol-seeking behavior, excessive alcohol consumption, a vulnerability to craving and relapse from alcohol-associated sensory cues, and persisting alcohol use despite adverse consequences (Cardenas et al., 2018; Ghin et al., 2022). There has been a growing body of evidence indicating that aberrant functional connectivity contributes to dysregulation in multiple brain networks in AUD (Kinreich, McCutcheon, et al., 2021; Meyers et al., 2021; Song et al., 2024). Several studies employing fMRI have found AUD associations with hypo-connectivity in frontoparietal executive control networks and hyper-connectivity in salience brain networks compared to healthy controls (Kamarajan et al., 2020; Suk et al., 2021; Weiland et al., 2014). Similarly weak resting state connectivity in left executive control networks has also been reported in opioid use disorder, indicating that this pattern of hypo-connectivity in executive control networks could be a general feature of substance use disorders (Woisard et al., 2021). Numerous investigations into the relationship between AUD on EEG coherence, have reported significantly greater coherence in the theta frequency band compared to non-alcoholic controls indicating aberrant hyper-connectivity between brain hemispheres (De Bruin et al., 2004; Kim et al., 2020; Meyers et al., 2021; Porjesz & Rangaswamy, 2007). The relationship between these patterns of hyper- and hypo-connectivity and AUD symptomatology remains unclear.

It is generally understood that the oscillatory dynamics of the brain reflect the coordinated interactions between groups of neurons across multiple temporal and spatial scales (Buzsáki, 2006; van Bree et al., 2025). While EEG coherence represents a within-frequency measures of functional connectivity, PAC represents cross-frequency connectivity dynamics which also take place within and between brain networks (Canolty & Knight, 2010). Cross-frequency coupling (CFC) is an umbrella term referring to putative mechanisms for coordinating the activity of neural assemblies oscillating at different frequencies, and these mechanisms have been associated with both healthy functioning as well as psychiatric and neurological disorders (Yakubov et al., 2022). Phase-amplitude coupling (PAC) is one means by which functional connectivity can be achieved where fast, high amplitude oscillations in one neuronal network are preferentially generated at a phase angle, often at peak or trough, of a slower frequency oscillation from another network (Deshpande et al., 2022; Huber et al., 2021; Marzetti et al., 2019). Statistical estimates of both PAC and coherence have minimum and maximum possible values, with EEG coherence ranging between the absence of synchrony to perfect synchronization, and PAC strength ranging from the absence of any modulatory relationship between the phase of one frequency to the amplitudes at a higher frequency (no coupling) to perfect coupling.

Comparisons of PAC strength between two conditions or with a control baseline, similarly to EEG coherence, are often interpreted as falling along an axis of hypo-connectivity to hyper-connectivity (Gonzalez et al., 2016; Hu et al., 2010; Leocani & Comi, 1999). For example, reduced PAC strength has been associated with diminished emotional face discrimination by those with autism spectrum disorder (Khan et al., 2013), lower cognitive functioning scores in patients with mild cognitive impairment (Musaeus et al., 2020), and poor sleep quality in individuals with Insomnia disorder (Guo et al., 2023), while excessive PAC magnitude has been associated with pathological symptoms in obsessive compulsive disorder (Bahramisharif et al., 2016; Treu et al., 2021; Yakubov et al., 2022), and in Parkinson’s disease (de Hemptinne et al., 2013; López-Azcárate et al., 2010; van Wijk et al., 2016; Yin et al., 2022). As research into cross frequency coupling in the brain has expanded over the past two decades, this initial interpretation has given way to more sophisticated models where changes in PAC strength have different effects depending on the neural circuits where they appear and in the context of within-frequency and cross-frequency manifestations of neural synchrony (Gong et al., 2022; Stujenske et al., 2014).

Many of the studies investigating putative roles that PAC plays in addictive disorders have centered on drug effects on the medial prefrontal cortex (mPFC), a core part of the brain reward system that has been shown to be intimately involved with alcohol and drug-related learning and memory (Abernathy et al., 2010; Sun et al., 2011). Increased theta-gamma PAC strength within prelimbic cortex, a part of mPFC, has been demonstrated in heroin addicted rats during conditioned place preference (Z. Zhu et al., 2019). Rats trained to associate cocaine delivery with foot shock persisted in cocaine seeking behavior after inactivation of prelimbic cortex by GABA agonists highlighting the role this part of the mPFC might play in terminating motivated behaviors that are harmful or noxious (Limpens et al., 2015). Moreover, long-term alcohol use appears to modify the mPFC in such a way as to facilitate impulsive or compulsive behaviors, comorbid features of AUD (Klenowski & 2018). The mPFC appears to encode the intention to drink alcohol, and when the contribution of mPFC in decision-making to drink alcohol is diminished, subcortical brain regions involved in habitual behaviors are possibly left as the primary driver of alcohol consumption (Linsenbardt et al., 2019).

Identifying new measures to characterize brain networks and functions underlying AUD will deepen our understanding of risk factors and resilience and enable designing effective treatments for AUD. While aberrant functional connectivity in the brain has been seen with fMRI and EEG coherence in AUD, the potential impacts of AUD on cross-frequency forms of functional connectivity like PAC have not yet been investigated.

In this project, we sought to determine whether PAC significantly differs between AUD and unaffected control groups, and if so, to confirm and characterize the phase-amplitude (PA) frequency pair domains where significant PAC differences were found. Our understanding of the neurophysiological, genetic, and environmental underpinnings of AUD has benefited greatly over the past several decades from large scale family-based datasets, like the Collaborative Study on the Genetics of Alcoholism (COGA), collecting multi-modal longitudinal data from families afflicted by AUD (Agrawal et al., 2023; Dick et al., 2023; Johnson et al., 2023). EEG resting state data drawn from COGA participants was used to compare resting state PAC estimates between those with severe AUD (DSM-5, 6+ symptoms) and unaffected controls. We focused on the frontomedial EEG channel FZ to illuminate any possible associations between AUD and PAC in the mPFC.

## METHODS

### Sample

The sample in the current study was drawn from the Collaborative Study on the Genetics of Alcoholism (COGA), a decades-long longitudinal national project following families affected with AUD and comparison families in a carefully characterized enriched family sample with data collection in multiple domains to investigate biological mechanisms underlying risk, development and consequences of alcohol addiction (Agrawal et al., 2023; Ehringer, 2023; Johnson et al., 2023; Kinreich, Meyers, et al., 2021; Meyers et al., 2023).

COGA participants completed comprehensive diagnostic clinical interviews (SSAGA) from which documentation about their past and present status of alcohol-related symptoms and diagnosis was gathered and used to determine whether they satisfied DSM-5 criteria for AUD (APA, 2013; Bucholz et al., 1994; Dick et al., 2023), along with a battery of EEG assessments, including resting state recordings (Meyers et al., 2023). Participants were told to arrive sober for the EEG recording session and they were rescheduled if they had a positive breath alcohol test. In this study, associations between AUD diagnosis and PAC were assessed using EEG recordings acquired during the resting state eyes closed condition as part of the COGA project. Detailed information on EEG collection in COGA is available in our previous publications (Begleiter et al., 1995; Meyers et al., 2023; Rangaswamy et al., 2002).

COGA participants between 25 and 50 years of age were selected to maximize the likelihood that results from this sample would reflect the typical adult brain without potential confounds in interpretation from developmental or aging processes, as the human brain, and the prefrontal cortex in particular, does not reach full maturity to adulthood until the mid-to-late 20s (Arain et al., 2013).

Control and experimental groups were age-matched for further control over possible age-related confounds (Table 1). Many of the COGA families were densely affected with AUD, so that COGA participants, including unaffected individuals used as controls in this study, were more likely to come from families at higher risk for AUD than the general population.

**Table 1.**
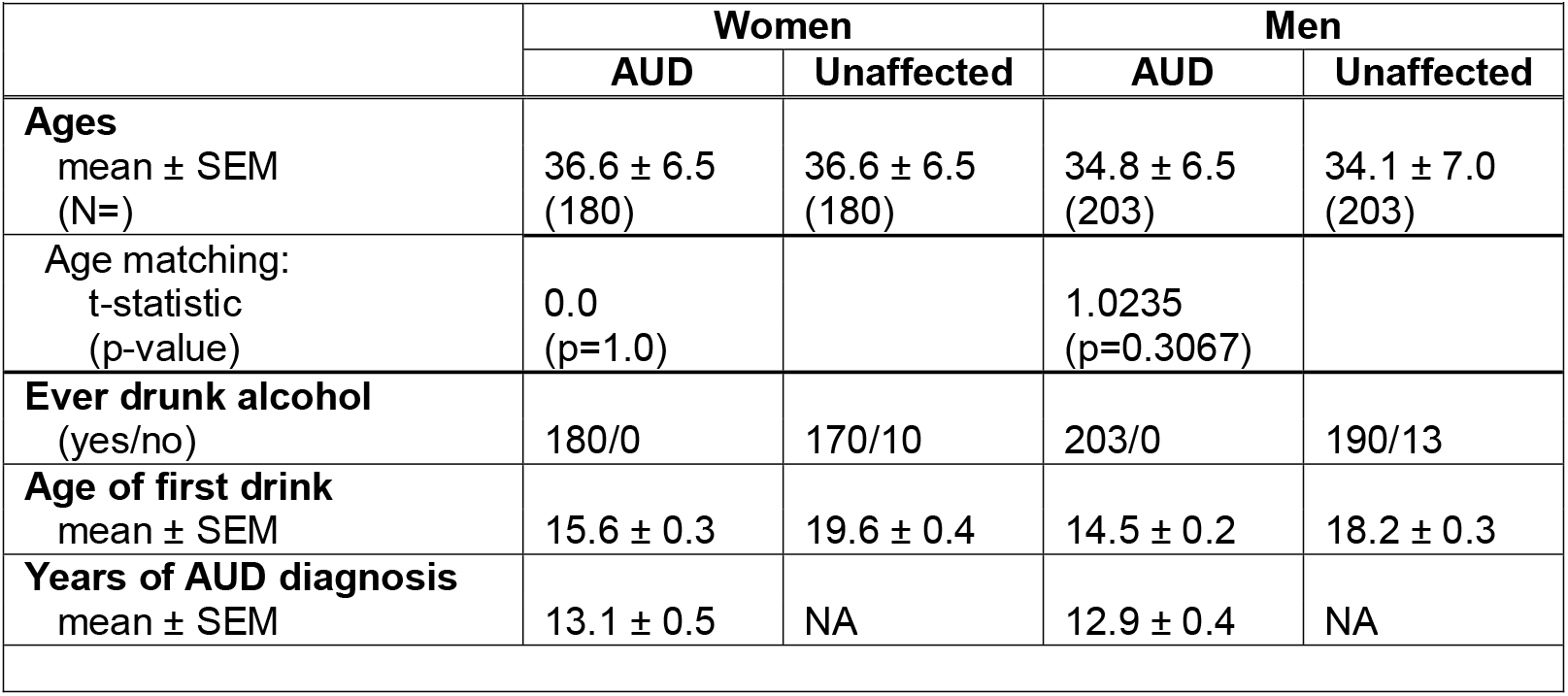
Participant demographics.

| Table 1. Participant demographics |  |  |  |  |
| --- | --- | --- | --- | --- |
|  | Women |  | Men |  |
|  | AUD | Unaffected | AUD | Unaffected |
| **Ages**<br>mean ± SEM<br>(N=) | 36.6 ± 6.5<br>(180) | 36.6 ± 6.5<br>(180) | 34.8 ± 6.5<br>(203) | 34.1 ± 7.0<br>(203) |
| Age matching:<br>t-statistic<br>(p-value) | 0.0<br>(p=1.0) |  | 1.0235<br>(p=0.3067) |  |
| **Ever drunk alcohol**<br>(yes/no) | 180/0 | 170/10 | 203/0 | 190/13 |
| **Age of first drink**<br>mean ± SEM | 15.6 ± 0.3 | 19.6 ± 0.4 | 14.5 ± 0.2 | 18.2 ± 0.3 |
| **Years of AUD diagnosis**<br>mean ± SEM | 13.1 ± 0.5 | NA | 12.9 ± 0.4 | NA |

We compared unaffected participants with those diagnosed with severe AUD, defined as having 6 or more of the 11 criteria for AUD diagnosis (DSM-5), to increase the signal-to-noise ratio and mitigate the possibility that PAC measures for unaffected COGA participants share greater similarities with AUD participants than those unaffected with no family history of AUD.

Overall, 766 subjects (N = 406 men) were included in this study (Table 1). Analyses were stratified by sex due to previous studies showing differences in brain structure and function between the sexes.

### EEG acquisition and preprocessing

Participants were fitted with a multi-channel electrode cap that included channel FZ in the EEG montage, with electrodes positioned at the nose as reference, and forehead as ground. During the acquisition of resting state EEG data, participants were seated in low lighting conditions in a temperature-controlled, sound-attenuated booth, and given instructions to remain awake and relaxed with eyes closed for the duration of the 5-minute session. The sampling rate was 256 Hz or 512 Hz depending on when COGA project data was collected. Impedances were kept below 5 kΩ for all EEG recordings.

Preprocessing of raw EEG signals included a high pass (1 Hz) and low pass (100 Hz) filtering with a zero-phase notch filter to remove the 60 Hz line noise. EEG signals were re-referenced to the average across EEG montage to permit use of MNE-ICAlabel package that identifies and removes non-brain artifacts (Li et al., 2022). Independent components analysis was performed using infomax method generating 15 independent components for use in MNE-ICAlabel artifact processing. Only independent components classified as ‘brain’ or ‘other’ were retained for signal reconstruction, while those labeled as ‘artifacts’ were removed. Python package MNE was used for EEG preprocessing steps (Gramfort et al., 2013). Coding infrastructure for this project which includes EEG preprocessing, PAC comodulogram image generation, and statistical analyses was built using Python and is available at https://github.com/lifepupil.

### Phase-Amplitude Coupling

PAC was estimated in EEG recordings from midfrontal channel FZ for every 30 seconds of EEG data. The 30 second EEG segments were decomposed into time-frequency representations in real and imaginary planes by wavelet transform using complex Morlet wavelet with number of cycles parameter in Gaussian width set to 7. PAC estimates were calculated for phase frequencies between 0.1-13 Hz, and amplitude frequencies between 4-50 Hz using 1 Hz window width at 0.1 Hz step to generate high resolution comodulograms.

At each 0.1 Hz step, the time-series of instantaneous amplitudes *φ*_*A*_ at frequency F_A_ and of instantaneous phase *φ*_*P*_ at frequency F_P_ were extracted from real and imaginary time-frequency representations of the EEG segment to calculate raw PAC estimates for the phase-amplitude frequency pair (F_P_, F_A_) using the Phase-Locking Value (PLV) equation

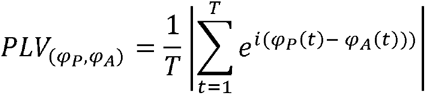

which measures consistency of phase differences between the phase *φ*_*P*_ and the phase of the amplitude envelope *φ*_*A*_ for all time points T in the signal (Lachaux et al., 1999; Seymour et al., 2017). A null distribution was generated by swapping amplitude time blocks prior to PAC estimation (200 permutations) to get expected mean value µ_0_ and standard deviation σ_0_ of PAC obtained by chance. Corrected PAC estimates were calculated by subtracting µ_0_ from the raw PAC estimates then dividing by σ_0_. Phase-amplitude comodulograms depicting the corrected PAC estimates across the phase frequency (x-axis) by amplitude frequency (y-axis) ranges were saved as grayscale images in JPG format and resized to 224 x 224 resolution (8-bit, 96 dpi) for use in a machine learning project described elsewhere (Richard et al., 2024). Normalization of PAC estimates from comodulograms was accomplished by rescaling 8-bit values from range [0,255] to [0,1]. The average comodulogram for each participant was calculated for the first two minutes of the resting state task (i.e., the first 4 comodulograms) to maximize the likelihood that the resting state EEG data used for analysis was gathered when participants were relaxed and awake with eyes closed but not drowsy or asleep. Statistical analyses were conducted on the resulting set of averaged comodulograms. Computational demands of PAC processing limited our analysis to a single EEG channel with FZ selected to target activity in the mPFC. TensorPAC package was used for all stages of PAC processing (Combrisson et al., 2020).

### PAC Comodulogram Statistical Analysis

PAC estimates from AUD and unaffected participants were compared at each phase-amplitude frequency pair (F_P_, F_A_) represented in the comodulogram using Mann-Whitney U test (two-tailed, α = 0.05). Test results were written to a set of matrices structured with same dimensions as the comodulograms. Resulting p-values from the test at (F_P_, F_A_) were stored in the comodulogram significance matrix (S-matrix) at the corresponding position S(F_P_, F_A_). Similarly, the differences between group PAC medians at (F_P_, F_A_), with unaffected group subtracted from the AUD group, were stored at D(F_P_, F_A_) in comodulogram difference matrix (D-matrix). Comodulogram median matrices U and A for unaffected and AUD groups, respectively, were similarly generated with median PAC at (F_P_, F_A_) from unaffected group comodulograms stored at U(F_P_, F_A_), as was done for AUD group median stored at A(F_P_, F_A_). Men and women were analyzed separately.

Once all frequency pairs had been tested, the comodulogram significance matrix was inspected for regions where the tests were significant across many neighboring frequency pairs, or clusters. Rectangular boundaries for each candidate cluster were determined by positioning a window of analysis, i.e., a cluster-specific PA domain, to tightly encompass the cluster of significant p-values in the S-matrix and the maximum PAC differences in the cluster as it appears in the D-matrix. Inclusion criteria for the initial set of candidate clusters were imposed for both matrices. The adjacent significant p-values comprising the cluster in the S-matrix had to span at least 1 Hz along the axis of either the amplitude or phase frequency, and their accompanying cluster of PAC differences at it appeared in the D-matrix had to span at least 1 Hz x 1 Hz in size. Multiple small but closely neighboring clusters were grouped together if they all fell within a PA domain with dimensions at least 1 Hz x 1 Hz. Only clusters considered for analysis were those falling within one of the standard PA domains represented on the comodulograms analyzed in this study, which include the delta-gamma, theta-gamma, alpha-gamma, delta-beta, theta-beta, alpha-beta, delta-alpha, and theta-alpha domains. These domain boundaries are based on standard EEG frequency bands of delta (0.1-3 Hz), theta (3-8 Hz), alpha (8-13 Hz), beta (13-28 Hz), and gamma (28-50 Hz).

A cluster p-value (P_FDR_) representing significance after correction for multiple comparisons was then calculated for each candidate by applying Benjamini-Hochberg on raw p-values within the cluster-specific PA domain of the S-matrix. Candidate clusters with P_FDR_ > 0.05 were excluded from further consideration. Cluster-specific comparisons of PAC by AUD diagnosis were conducted on the remaining candidates by calculating median PAC for each participant, then performing group comparisons of the cluster median PAC estimates with Mann-Whitney U test (two-tailed, α = 0.05). Cluster PAC strength for each participant was calculated by taking the median PAC from within the boundaries of the cluster-specific PA domain from that participant’s average comodulogram (described in previous section). Only candidate clusters satisfying all validation and statistical testing conditions were reported as significant PAC clusters.

Measures of effect size and correlation appropriate to nonparametric testing were calculated for each candidate cluster. The Common language effect size (CLES) indicates the probability of PAC being greater in someone randomly selected from the AUD group compared to someone randomly selected from the unaffected group. The rank biserial correlation coefficient, r, represents the magnitude and direction of the relationship between AUD diagnosis and ordinal dependent variable of ranked PAC strength. This coefficient has a range [-1,1], where a perfect positive relationship (r=1) would be obtained if ranks from AUD group are all higher than those from the unaffected group. Conversely, a perfect negative relationship (r=-1) would result if all ranks from AUD group were lower than any from the unaffected group. In both cases, the rank distributions of the two groups are non-overlapping. A coefficient r=0 indicates perfect overlap of the two rank distributions, i.e., where there are no differences between the distribution of ranks for AUD and unaffected groups.

## RESULTS

Review of comodulogram significance matrix revealed clusters qualifying for further investigation in both men and women (Figure 1A). The initial set of candidate clusters (17 for women; 17 for men) were distributed across all standard PA domains represented in the comodulograms, though not uniformly so (Figure 1A, green lines indicate standard PA domain boundaries). Among women, apart from the large candidate clusters in alpha-gamma domain, most candidate clusters were concentrated in PA domains encompassing delta/low theta (0.1-5 Hz) phase frequency bands paired with alpha/low beta (8-15 Hz) or high beta/gamma (20-50 Hz) amplitude frequency bands, most conspicuously within delta-alpha, theta-alpha, and delta-gamma domains (Figure 1A). Like women, apart from large candidate clusters in alpha-gamma, over half of the candidates identified in men were found in PA domains encompassing delta/low theta (0.1-5 Hz) phase frequency bands including large candidate clusters in the theta-gamma domain, with the rest appearing in alpha-beta and neighboring theta-beta domains.

**Figure 1.**
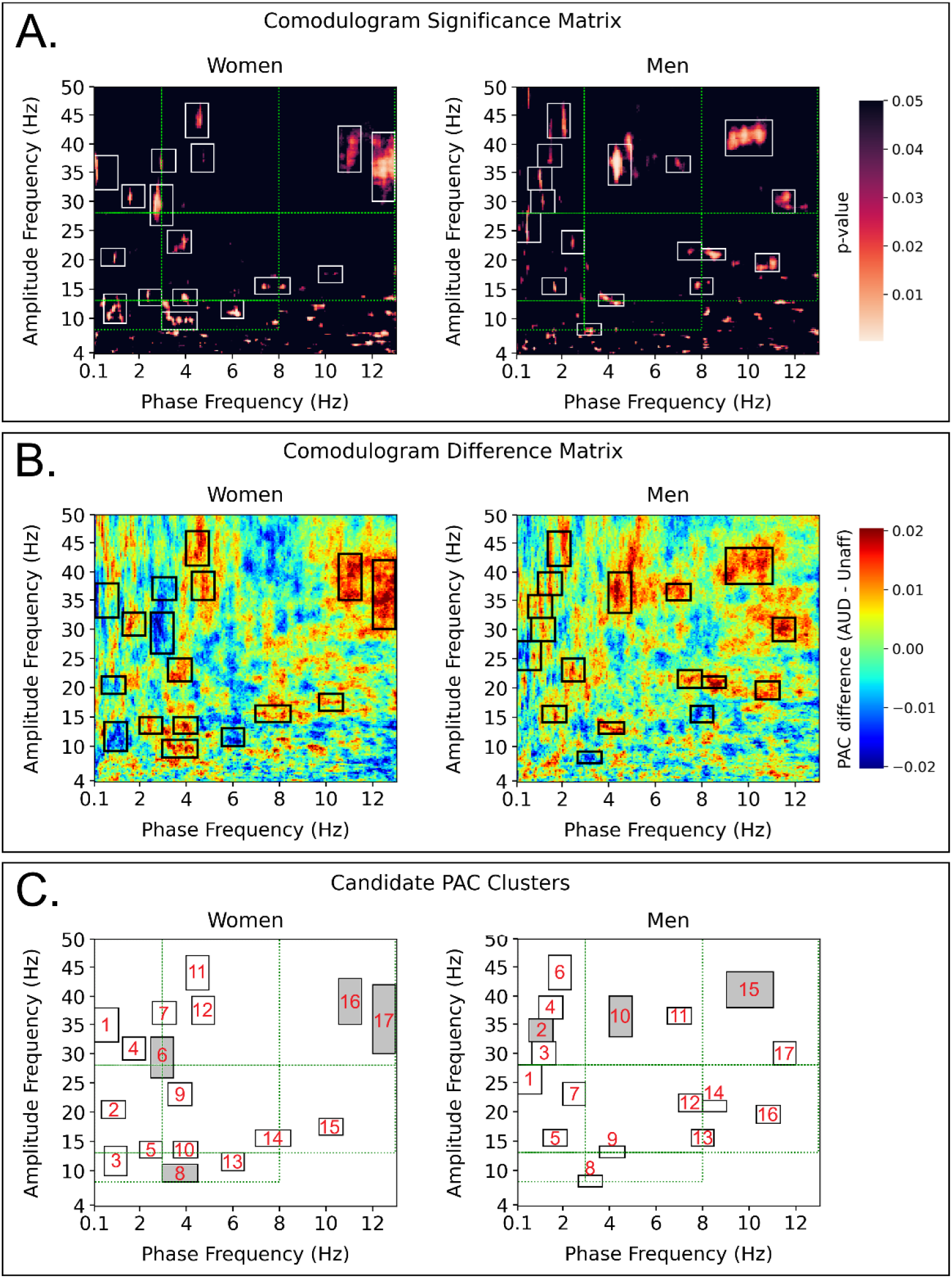
Difference and significance matrix results from comodulogram analysis. [A]: comodulogram significance matrix (S-matrix) for women (left) and men (right) masked to only reveal p-values less than or equal to 0.05. PA domains of candidate clusters marked with black rectangles. PAC domains outlined in green for reference. [B]: comodulogram difference matrix results (D-matrix) by sex. [C]: PA domains of candidate clusters by sex. Cluster boundaries are marked with black rectangles. Numbering corresponds to the cluster ID in Table 2.

With respect to group differences in PAC strength, in men, most candidate clusters in comodulogram difference matrix showed greater PAC in the AUD group relative to unaffected, particularly in delta-gamma, theta-gamma, and alpha-gamma domains (Figure 1B) with the remaining candidate clusters appearing across beta frequency ranges of the amplitude frequency, e.g., in the delta-beta, theta-beta, and alpha-beta domains. In women, too, PAC strength of candidate clusters in theta-gamma and alpha-gamma domains was generally greater in AUD than in the unaffected group, though 3 of the 4 candidate clusters in delta-gamma domain showed lower PAC in the AUD group (Figure 1B). Two candidate clusters were found with almost identical PA domains of both men and women, including women’s cluster 8 (W8) with men’s cluster 9 (M9) on the boundary of theta-alpha and theta-beta domains, and W14 with M13 on the boundary of theta-beta and alpha-beta domains (Figure 1C). While candidate clusters W8 and M9 showed greater PAC in the AUD group, PAC differences in candidate clusters W14 and M13 would appear to be sexually dimorphic in AUD groups with lower PAC in men and greater PAC in women. Candidate clusters were numbered in order of appearance along the phase frequency axis starting at 0.1 Hz (Figure 1C).

**Table 2.**
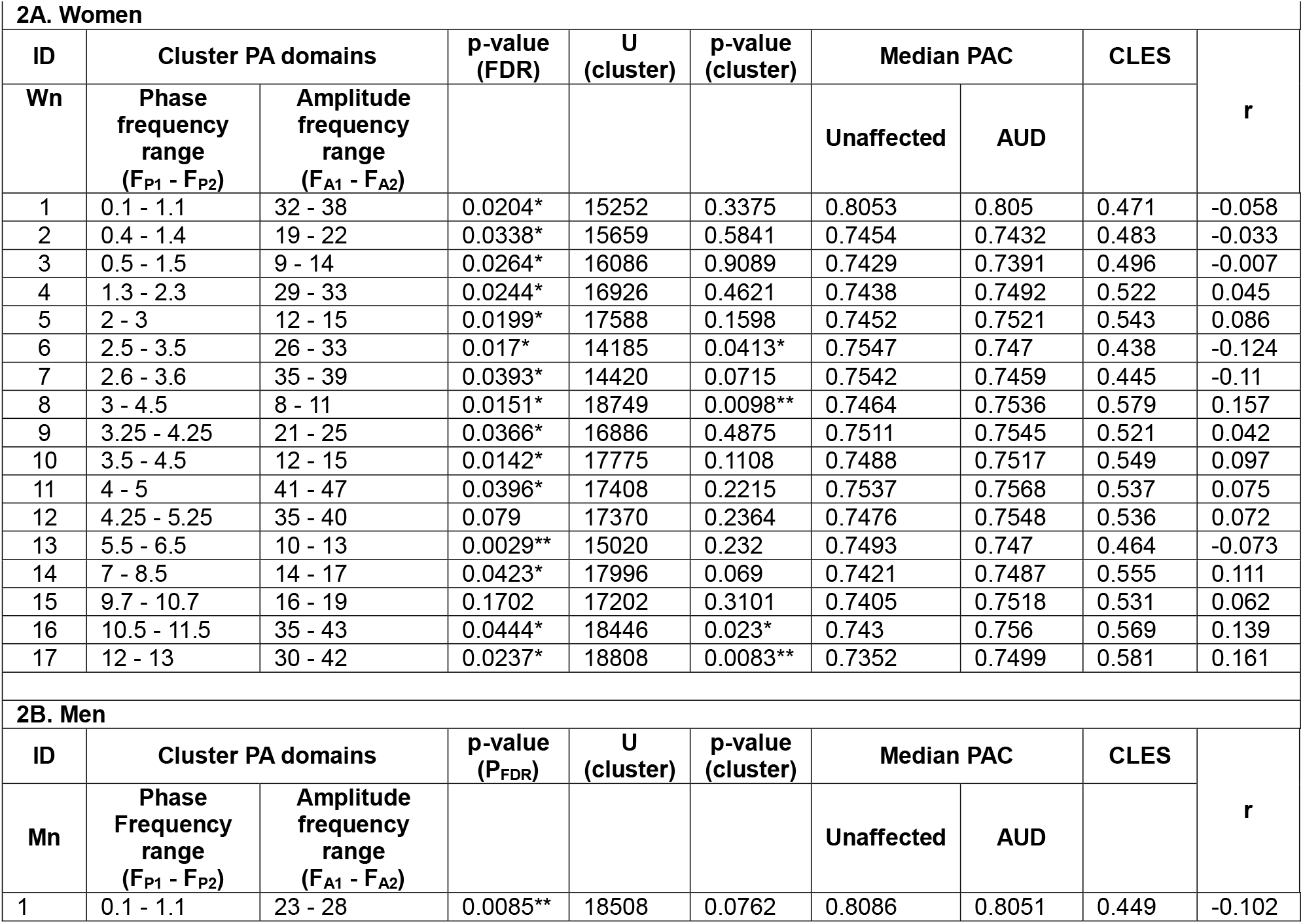
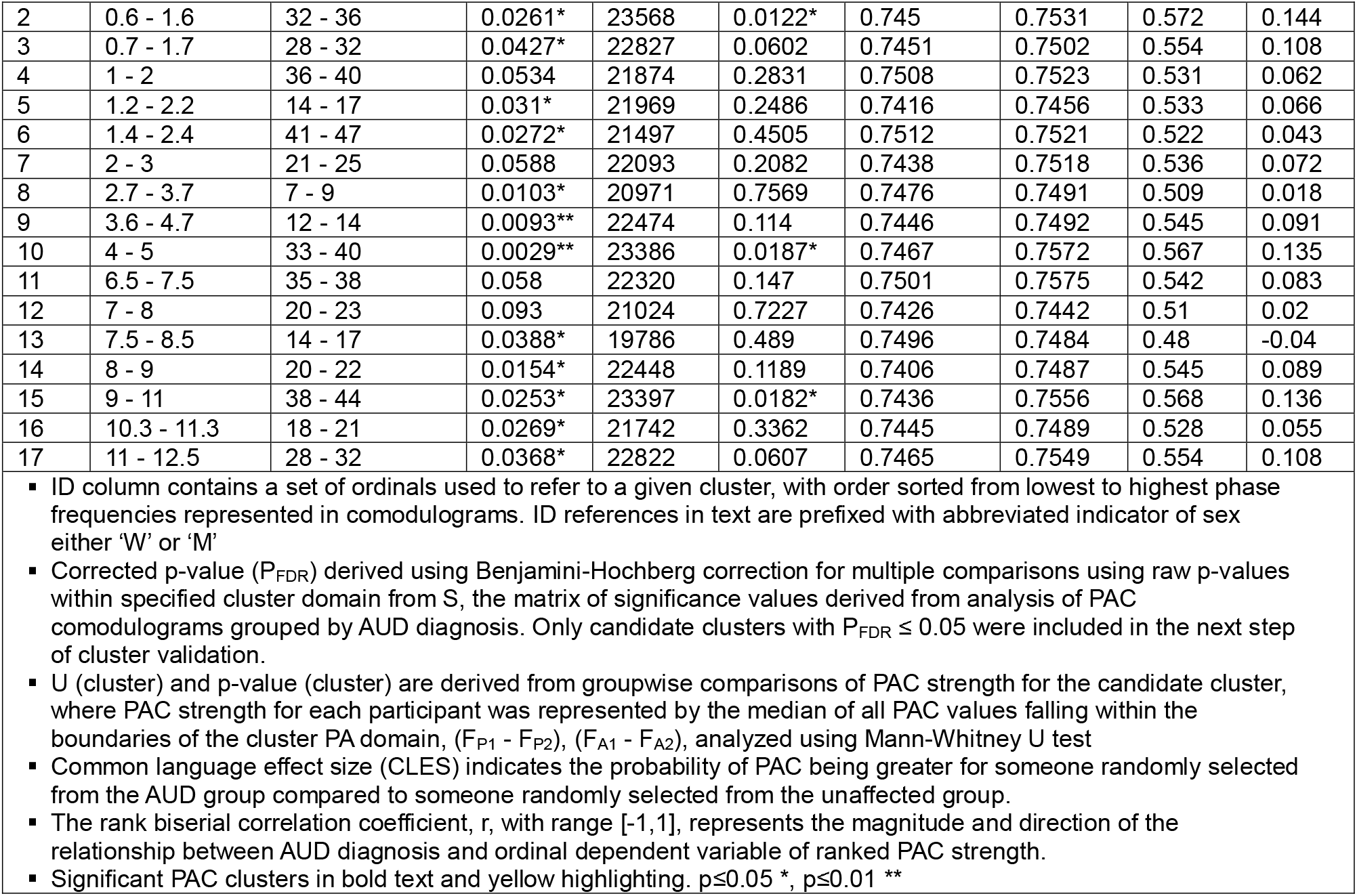
Phase-Amplitude Coupling Cluster Analysis.

| **2A. Women** |  |  |  |  |  |  |  |  |  |
| --- | --- | --- | --- | --- | --- | --- | --- | --- | --- |
| ID | Cluster PA domains |  | p-value (FDR) | U (cluster) | p-value (cluster) | Median PAC |  | CLES | r |
| Wn | Phase frequency range (F<sub>P1</sub> - F<sub>P2</sub>) | Amplitude frequency range (F<sub>A1</sub> - F<sub>A2</sub>) |  |  |  | Unaffected | AUD |  |  |
| 1 | 0.1 - 1.1 | 32 - 38 | 0.0204* | 15252 | 0.3375 | 0.8053 | 0.805 | 0.471 | -0.058 |
| 2 | 0.4 - 1.4 | 19 - 22 | 0.0338* | 15659 | 0.5841 | 0.7454 | 0.7432 | 0.483 | -0.033 |
| 3 | 0.5 - 1.5 | 9 - 14 | 0.0264* | 16086 | 0.9089 | 0.7429 | 0.7391 | 0.496 | -0.007 |
| 4 | 1.3 - 2.3 | 29 - 33 | 0.0244* | 16926 | 0.4621 | 0.7438 | 0.7492 | 0.522 | 0.045 |
| 5 | 2 - 3 | 12 - 15 | 0.0199* | 17588 | 0.1598 | 0.7452 | 0.7521 | 0.543 | 0.086 |
| 6 | 2.5 - 3.5 | 26 - 33 | 0.017* | 14185 | 0.0413* | 0.7547 | 0.747 | 0.438 | -0.124 |
| 7 | 2.6 - 3.6 | 35 - 39 | 0.0393* | 14420 | 0.0715 | 0.7542 | 0.7459 | 0.445 | -0.11 |
| 8 | 3 - 4.5 | 8 - 11 | 0.0151* | 18749 | 0.0098** | 0.7464 | 0.7536 | 0.579 | 0.157 |
| 9 | 3.25 - 4.25 | 21 - 25 | 0.0366* | 16886 | 0.4875 | 0.7511 | 0.7545 | 0.521 | 0.042 |
| 10 | 3.5 - 4.5 | 12 - 15 | 0.0142* | 17775 | 0.1108 | 0.7488 | 0.7517 | 0.549 | 0.097 |
| 11 | 4 - 5 | 41 - 47 | 0.0396* | 17408 | 0.2215 | 0.7537 | 0.7568 | 0.537 | 0.075 |
| 12 | 4.25 - 5.25 | 35 - 40 | 0.079 | 17370 | 0.2364 | 0.7476 | 0.7548 | 0.536 | 0.072 |
| 13 | 5.5 - 6.5 | 10 - 13 | 0.0029** | 15020 | 0.232 | 0.7493 | 0.747 | 0.464 | -0.073 |
| 14 | 7 - 8.5 | 14 - 17 | 0.0423* | 17996 | 0.069 | 0.7421 | 0.7487 | 0.555 | 0.111 |
| 15 | 9.7 - 10.7 | 16 - 19 | 0.1702 | 17202 | 0.3101 | 0.7405 | 0.7518 | 0.531 | 0.062 |
| 16 | 10.5 - 11.5 | 35 - 43 | 0.0444* | 18446 | 0.023* | 0.743 | 0.756 | 0.569 | 0.139 |
| 17 | 12 - 13 | 30 - 42 | 0.0237* | 18808 | 0.0083** | 0.7352 | 0.7499 | 0.581 | 0.161 |
| **2B. Men** |  |  |  |  |  |  |  |  |  |
| ID | Cluster PA domains |  | p-value (P<sub>FDR</sub>) | U (cluster) | p-value (cluster) | Median PAC |  | CLES | r |
| Mn | Phase Frequency range (F<sub>P1</sub> - F<sub>P2</sub>) | Amplitude frequency range (F<sub>A1</sub> - F<sub>A2</sub>) |  |  |  | Unaffected | AUD |  |  |
| 1 | 0.1 - 1.1 | 23 - 28 | 0.0085** | 18508 | 0.0762 | 0.8086 | 0.8051 | 0.449 | -0.102 |
| 2 | 0.6 - 1.6 | 32 - 36 | 0.0261* | 23568 | 0.0122* | 0.745 | 0.7531 | 0.572 | 0.144 |
| 3 | 0.7 - 1.7 | 28 - 32 | 0.0427* | 22827 | 0.0602 | 0.7451 | 0.7502 | 0.554 | 0.108 |
| 4 | 1 - 2 | 36 - 40 | 0.0534 | 21874 | 0.2831 | 0.7508 | 0.7523 | 0.531 | 0.062 |
| 5 | 1.2 - 2.2 | 14 - 17 | 0.031* | 21969 | 0.2486 | 0.7416 | 0.7456 | 0.533 | 0.066 |
| 6 | 1.4 - 2.4 | 41 - 47 | 0.0272* | 21497 | 0.4505 | 0.7512 | 0.7521 | 0.522 | 0.043 |
| 7 | 2 - 3 | 21 - 25 | 0.0588 | 22093 | 0.2082 | 0.7438 | 0.7518 | 0.536 | 0.072 |
| 8 | 2.7 - 3.7 | 7 - 9 | 0.0103* | 20971 | 0.7569 | 0.7476 | 0.7491 | 0.509 | 0.018 |
| 9 | 3.6 - 4.7 | 12 - 14 | 0.0093** | 22474 | 0.114 | 0.7446 | 0.7492 | 0.545 | 0.091 |
| 10 | 4 - 5 | 33 - 40 | 0.0029** | 23386 | 0.0187* | 0.7467 | 0.7572 | 0.567 | 0.135 |
| 11 | 6.5 - 7.5 | 35 - 38 | 0.058 | 22320 | 0.147 | 0.7501 | 0.7575 | 0.542 | 0.083 |
| 12 | 7 - 8 | 20 - 23 | 0.093 | 21024 | 0.7227 | 0.7426 | 0.7442 | 0.51 | 0.02 |
| 13 | 7.5 - 8.5 | 14 - 17 | 0.0388* | 19786 | 0.489 | 0.7496 | 0.7484 | 0.48 | -0.04 |
| 14 | 8 - 9 | 20 - 22 | 0.0154* | 22448 | 0.1189 | 0.7406 | 0.7487 | 0.545 | 0.089 |
| 15 | 9 - 11 | 38 - 44 | 0.0253* | 23397 | 0.0182* | 0.7436 | 0.7556 | 0.568 | 0.136 |
| 16 | 10.3 - 11.3 | 18 - 21 | 0.0269* | 21742 | 0.3362 | 0.7445 | 0.7489 | 0.528 | 0.055 |
| 17 | 11 - 12.5 | 28 - 32 | 0.0368* | 22822 | 0.0607 | 0.7465 | 0.7549 | 0.554 | 0.108 |

- ID column contains a set of ordinals used to refer to a given cluster, with order sorted from lowest to highest phase frequencies represented in comodulograms. ID references in text are prefixed with abbreviated indicator of sex either 'W' or 'M' - Corrected p-value ( $P_{FDR}$ ) derived using Benjamini-Hochberg correction for multiple comparisons using raw p-values within specified cluster domain from S, the matrix of significance values derived from analysis of PAC comodulograms grouped by AUD diagnosis. Only candidate clusters with $P_{FDR} \leq 0.05$ were included in the next step of cluster validation. - U (cluster) and p-value (cluster) are derived from groupwise comparisons of PAC strength for the candidate cluster, where PAC strength for each participant was represented by the median of all PAC values falling within the boundaries of the cluster PA domain, ( $F_{P1} - F_{P2}$ ), ( $F_{A1} - F_{A2}$ ), analyzed using Mann-Whitney U test - Common language effect size (CLES) indicates the probability of PAC being greater for someone randomly selected from the AUD group compared to someone randomly selected from the unaffected group. - The rank biserial correlation coefficient, r, with range [-1,1], represents the magnitude and direction of the relationship between AUD diagnosis and ordinal dependent variable of ranked PAC strength. - Significant PAC clusters in bold text and yellow highlighting. $p \leq 0.05$ \*, $p \leq 0.01$ \*\*

Among the candidate clusters analyzed, in women, four were found to have satisfied validation and statistical testing conditions that reveal significant differences in PAC between AUD and unaffected groups (Table 2A), while in men, three significant PAC clusters were found of the 17 candidates tested (Table 2B).

The largest significant PAC clusters were found in the alpha-gamma domain of both men (M15, Table 2B) and women (W16, W17, Table 2A). The AUD group exhibited greater PAC than the unaffected group in all significant alpha-gamma clusters regardless of sex. Men in the AUD group had significantly greater PAC than the unaffected group in the M15 cluster (U=23397, p=0.0182, CLES=0.568, r = 0.136). The AUD group distribution of PAC estimates from this cluster revealed a significant shift towards greater PAC compared to the unaffected group distribution (Figure 2B, cluster 15: AUD median = 0.7556, unaffected median = 0.7436).

**Figure 2.**
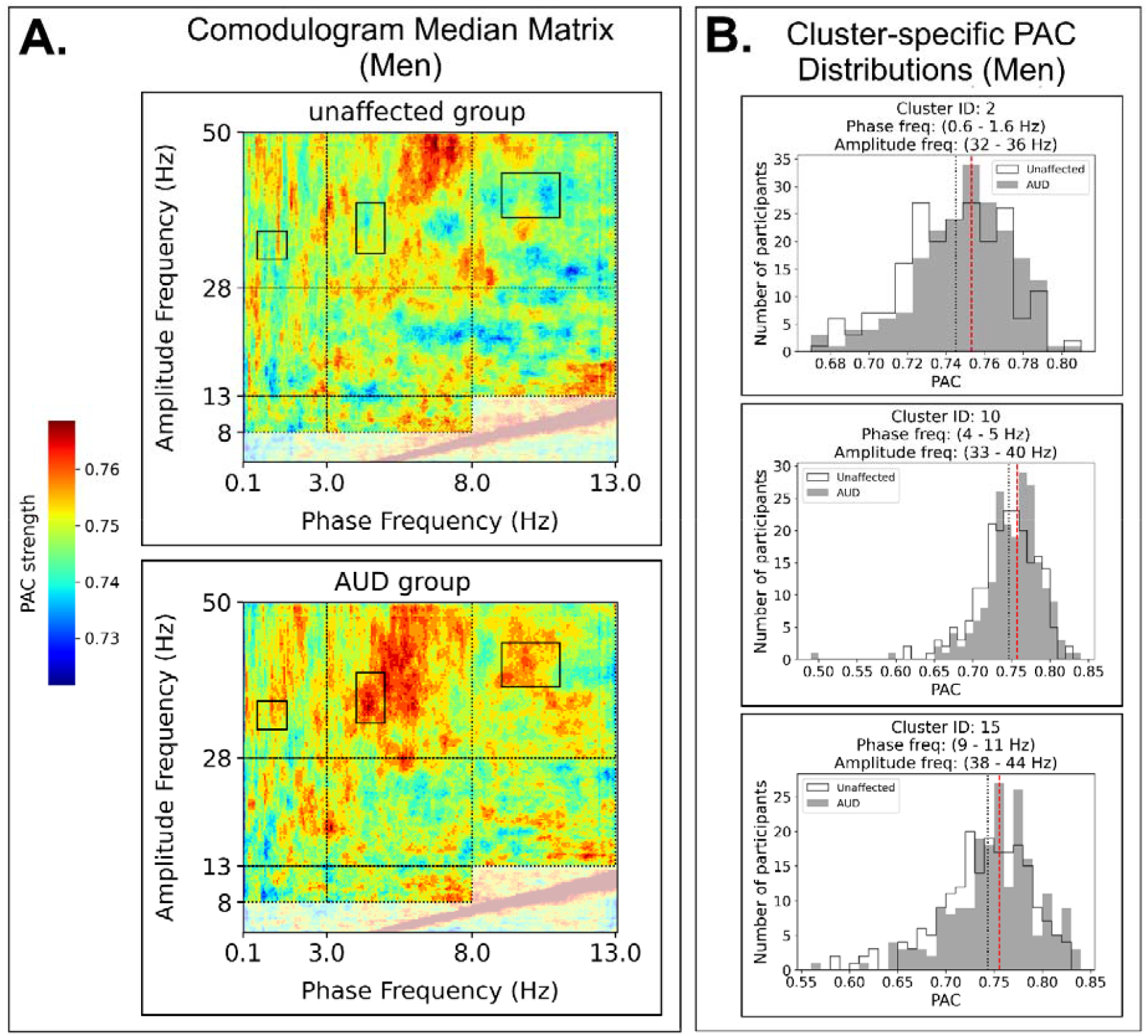
PA domains containing significant PAC clusters in men. Standard PA domain boundaries marked by black dotted lines. Only clusters considered for analysis were those falling within one of the standard PA domains represented on the comodulograms analyzed in this study, which include the delta-gamma, theta-gamma, alpha-gamma, delta-beta, theta-beta, alpha-beta, delta-alpha, and theta-alpha domains. These domain boundaries are based on standard EEG frequency bands of delta (0.1-3 Hz), theta (3-8 Hz), alpha (8-13 Hz), beta (13-28 Hz), and gamma (28-50 Hz). Faded regions of the comodulogram have been excluded from cluster analysis.

Among women, PAC was significantly greater for the AUD group compared to unaffected participants in alpha-gamma cluster W16 (U=18446, p=0.023, CLES=0.569, r = 0.139), and cluster W17 (U=18808, p=0.0083, CLES=0.581, r = 0.161). Like men with AUD, the distribution of PAC estimates from women in the AUD group were significantly shifted towards greater PAC compared to those in the unaffected group in clusters W16 (Figure 3B, cluster 16: AUD median = 0. 756, unaffected median = 0. 743) and W17 (Figure 3B, cluster 17: AUD median = 0.7499, unaffected median = 0.7352). Review of comodulogram median matrix results indicated that these significant PAC clusters were appearing in the context of greater PAC strength broadly across the alpha-gamma domain of both men (Figure 2A) and women (Figure 3A) with severe AUD compared to their unaffected counterparts.

**Figure 3.**
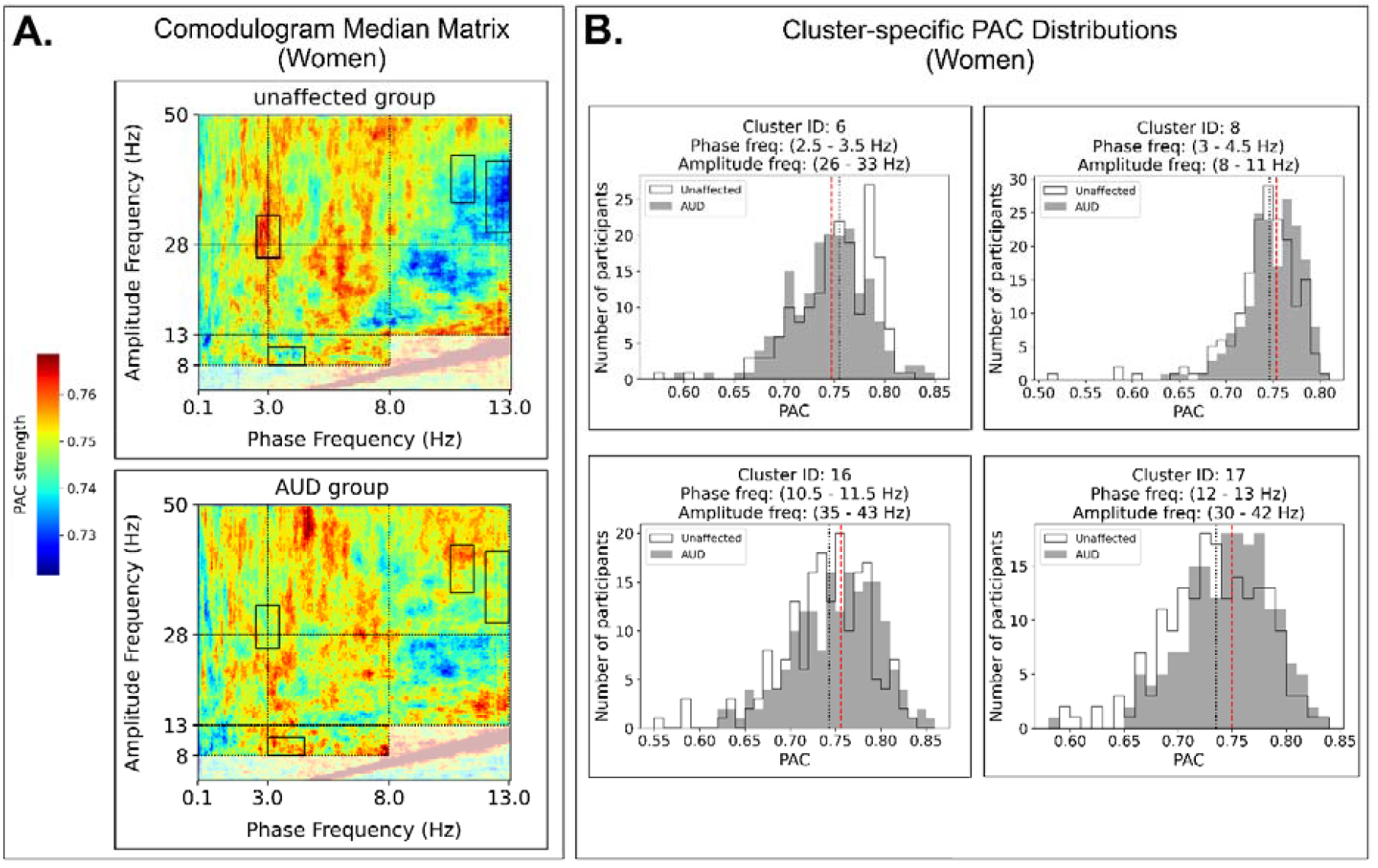
PA domains containing significant PAC clusters in women. Standard PA domain boundaries marked by black dotted lines. Only clusters considered for analysis were those falling within one of the standard PA domains represented on the comodulograms analyzed in this study, which include the delta-gamma, theta-gamma, alpha-gamma, delta-beta, theta-beta, alpha-beta, delta-alpha, and theta-alpha domains. These domain boundaries are based on standard EEG frequency bands of delta (0.1-3 Hz), theta (3-8 Hz), alpha (8-13 Hz), beta (13-28 Hz), and gamma (28-50 Hz). Faded regions of the comodulogram have been excluded from cluster analysis.

This pattern of broadly greater PAC strength in AUD groups appears to diverge by sex in the theta-gamma and delta-gamma domains (Figure 2A, Figure 3A). While candidate clusters in the theta-gamma domain indicated greater PAC in the AUD groups of both men (Figure 2A, cluster 10) and women (Figure 3A, clusters 11, 12), only the M10 cluster rose to significance (AUD median = 0.7572, unaffected median = 0.7467, U=23386, p=0.0187, CLES=0.567, r = 0.135).

Candidate clusters from both men and women were also found within the delta-gamma domain, albeit in mostly non-overlapping regions. The only significant PAC cluster seen in women (W6) straddles the boundaries of delta and theta bands (2.5 - 3.5 Hz) in phase frequency, and across beta/gamma boundaries (26 - 33 Hz) in amplitude frequency ranges. Cluster W6 is also notable in that it was the only cluster of significantly lower PAC in the AUD group compared to their unaffected counterparts (cluster 6: AUD median = 0.747, unaffected median = 0. 7547, U=23568, p=0.0122, CLES=0.572, r = 0.144).

Among men, significant PAC differences were found in the delta-gamma M2 cluster, with significantly greater PAC seen in men with AUD (cluster 2: AUD median = 0.7531, unaffected median = 0.745, U=23568, p=0.0122, CLES=0.572, r = 0.144). Compared to candidate and significant PAC clusters from the alpha-gamma and theta-gamma domains, those in the delta-gamma domain were mostly non-overlapping between the sexes. Only women were found to have a significant PAC cluster in the theta-alpha domain (cluster 8: AUD median = 0. 7536, unaffected median = 0. 7464, U=18749, p=0.0098, CLES=0.579, r = 0.157).

## DISCUSSION

The present study represents an initial assessment of cross-frequency functional connectivity in AUD by comparing PAC estimates from those with severe AUD to age-matched groups of unaffected participants. The primary objective was to identify PA domains that appear to be sensitive to AUD status to provide specific domains of interest to target in future studies and more generally characterize PAC activity recorded from frontomedial brain regions that include the mPFC.

We found evidence of PAC taking place in both AUD and unaffected groups under these conditions, albeit to varying degrees. Distributions of PAC estimates from all significant clusters ranged from roughly 0.6 to 0.85 in both men (Figure 2B) and women (Figure 3B) on a [0,1] scale which suggests that relatively small deviations in PAC might be diagnostically meaningful. A crossover study of alcohol intoxication in a sex-balanced group of healthy adults found small but significant reductions in delta-gamma and theta-gamma PAC to frontal and parietal EEG channels during resting state when under the influence of alcohol (Lee & Yun, 2014). The authors point out that the reduction in PAC strength is consistent with findings from other studies of alcohol intoxication-related decreased in functional connectivity, suggesting that the decreased PAC might reflect a disruption of information transfer between local neuronal assemblies and larger-scale brain networks. The reductions in delta-gamma and theta-gamma PAC during alcohol intoxication that they report contrast sharply with the greater PAC we saw, particularly among men with AUD. Though direct comparisons are not possible from the results of this study since we only have data acquired from participants while in a sober state, one wonders whether the AUD-associated increases in theta-gamma PAC might reflect compensatory neuroadaptations arising from years of repeated exposure to alcohol, though future studies will be needed to cast more light on these issues.

Cluster analysis conducted separately on men and women revealed apparent similarities and differences between the sexes. The greatest similarity between the sexes appeared in the alpha-gamma domain (Figure 2A, Figure 3A). All candidate clusters found in alpha-gamma domain showed greater PAC in the AUD groups, and significant alpha-gamma PAC clusters were seen in both men (M15, Figure 2B) and women (W16, W17, Figure 2B).

Similar frontal EEG resting state measures of excessive alpha-gamma PAC strength that we found in the severe AUD group has also been reported in obsessive-compulsive disorder (OCD) patients undergoing deep brain stimulation (Figee et al., 2013; Treu et al., 2021). Abnormally elevated functional connectivity of ventrolimbic corticostriatal regions in patients with OCD has been positively correlated with severity of symptoms (Harrison et al., 2009). Compared to controls, OCD patients exhibited significant increases in alpha-gamma and low beta-gamma PAC over frontal midline and parietal electrodes when deep brain stimulation (DBS) of nucleus accumbens (NAc) was turned off, and was reduced when NAc-DBS was active (Treu et al., 2021). In another study, researchers found that excessive beta-gamma PAC in frontostriatal circuits of OCD during resting state, eyes open was reduced with deep brain stimulation (DBS) of nucleus accumbens as was symptom severity (Figee et al., 2013).

Excessive resting state PAC has also been reported in primary motor cortex and adjacent motor regions of Parkinson’s disease (PD) patients in a PA domain encompassing the 10-25 Hz phase frequency by 30-200 Hz amplitude frequency range (de Hemptinne et al., 2013; López-Azcárate et al., 2010; van Wijk et al., 2016; Yin et al., 2022). Pathologically strong beta-gamma PAC seen in unmedicated Parkinson’s disease patients was associated with more severe motor symptoms, both of which were dramatically reduced following levodopa treatment (López-Azcárate et al., 2010; van Wijk et al., 2016). PD patients also exhibited strong beta-gamma PAC in the motor cortex accompanying freezing of gait with gait freezing temporally aligned with increased beta-gamma PAC (Yin et al., 2022). It has been suggested that observed excessive PAC might reflect a pathological restriction of motor cortex to a monotonous pattern of coupling that make it less responsive to signals from brain regions mediating voluntary movement (de Hemptinne et al., 2013). In fact, results from a recent study have provided evidence indicating that temporal PAC dynamics may be as important as changes in magnitude of PAC strength. EEG measured during a finger tapping task revealed a PAC motif unfolding across movement execution that was seen in both PD patients and controls, with bradykinesia-related deficits in the PD patients associated with aberrant expressions of this PAC motif (Gong et al., 2022). These results indicate the possibility that dynamic regulation of PAC across time might be at least as important as overall changes in PAC strength.

Clusters falling entirely within theta-gamma domain showed greater PAC in those with AUD diagnosis regardless of sex as can be seen in the overlapping cluster PA domains of W11, W12 and M10, indicating, at least in this sample, that sex differences in theta-gamma PAC are differences of magnitude and not of direction, though PAC differences were only significant in the theta-gamma clusters of men.

The increases in theta-gamma PAC strength we report here are consistent with the handful of previous studies reported for other substance abuse disorders. Notable among them are several animal models of addiction based on conditioned place preference (CPP). The CPP paradigm employs a test chamber divided into two sides separated by a wall; in the conditioning stage, animals are administered a drug or other reward prior to placement in one side of the chamber in which they are restricted, and no reward when placed on the opposite side. During the post-conditioning stage, animals are allowed unrestricted access, without reward treatment, to both sides of the chamber during which time spent on either side is measured (Fattahi et al., 2023). Animals spend more time on the side of the test chamber in which they had undergone repeated reward-pairing, having learned to associate the side-specific sensory stimuli with reward. In one such study, Zhu and colleagues gathered local field potentials (LFP) from the prelimbic area of mPFC in male rats before and after CPP induction with heroin (Z. M. Zhu et al., 2019). As expected, heroin-addicted rats spent significantly more time on the drug-paired side than the saline-paired side. During the post-conditioned stage, LFP revealed increased relative theta power accompanied by decreased relative gamma power, and significantly greater theta-gamma PAC in the same heroin-addicted rats. In another CPP animal study targeting the basolateral amygdala for LFP recordings (Nukitram et al., 2021), researchers found that methamphetamine-treated male mice exhibited significantly enhanced theta-gamma PAC during the post-conditioning stage compared to pre-conditioning baseline. This pattern of elevated theta-gamma PAC reported in these studies are not limited to drugs of abuse but has also been seen when testing place preference conditioned by highly palatable food (Samerphob et al., 2017), indicating that increases in theta-gamma PAC could be reflecting some aspect of reward-related sensory cues.

Differences between the sexes appeared to be greatest in the delta-gamma domain. Median delta-gamma PAC in men with AUD was broadly greater compared to unaffected men, as were all candidate clusters (M2, M3, M4, M6) and significantly greater in cluster M2. In contrast, most of the delta-gamma candidate clusters in women (W1, W6, W7) showed lower PAC in women with AUD compared to their unaffected counterparts, with significantly lower PAC found at the W6 cluster. One possible explanation for the lower PAC among women with AUD could be insufficient neural synchrony in the theta band, Lower theta EEG coherence (3-5 Hz) during resting state has been observed in women with AUD particularly at parietal brain regions (Meyers et al., 2021). If the effects of this reduced theta EEG coherence extend through parieto-frontal brain circuits, one might expect an accompanying alteration in PAC strength in recordings from frontal brain regions in the 3-5 Hz phase frequency range which would include the W6 cluster as well as W8 in the theta-alpha domain. Further investigation will be needed to test between this or other possible explanations.

## LIMITATIONS

The focus of the study on PAC activity within a single EEG channel does preclude any determination of the source of PAC, as either arising within local mPFC networks or driven by the activity of distal brain networks, that a multi-channel approach could deliver. However, the decision to restrict investigation to the frontomedial channel FZ represented a trade-off in service of the primary objective of the study, based on the absence of any other published studies on PAC in AUD, to identify any PA domains associated with AUD that could be targeted in follow-up investigations. Single channel analysis allowed us to reduce the considerable computing demands required to generate comodulograms for the 700+ participants represented in this study, providing greater confidence in the precision and accuracy of PAC estimates represented in the high resolution comodulogram data on which the cluster analyses were conducted.

## FUTURE DIRECTIONS

Replicate studies employing a multi-channel approach measuring PAC within the PA domains reported here across brain regions could both test whether similar results are found and to what extent these differences in PAC might appear in neighboring channels. This could include formal testing of sex differences at target PA domains between frontal and parietal EEG channels to determine whether these inter-channel PAC estimates might be contributing to the lower PAC among women with AUD. The results of this study open other possible lines of inquiry. The AUD-associated PA domains reported here were based on individuals with severe AUD, defined as having any 6 or more of the 11 AUD symptoms listed in the DSM-5, though it is unclear how individual symptoms might map onto these AUD-associated PA domains. It could be the case that there are PA domain-specific AUD symptoms, such as with physical dependence and withdrawal symptoms, or alcohol craving and seeking behaviors. Correlational analyses could be undertaken to test symptom-based hypotheses of PAC by PA domain, as well as for other cognitive or behavioral measures that appear to be affected by AUD status, comorbid features like impulsivity and compulsivity, or genomic association studies.

## ETHICS STATEMENT

All procedures involving human participants were performed in accordance with the ethical standards of the institutional research committee. This research was conducted at SUNY Downstate Health Sciences University under approved IRB protocol # 266920 (Ref #: 89-040, Date: 10/7/2025). This research used COGA data under approved IRB protocol # 20243210 (Date: 07/21/2025).

## ACKNOWLEDGMENTS

Support for this research came from the National Institute on Alcohol Abuse and Alcoholism (AA029448) and the Collaborative Study on the Genetics of Alcoholism project grant (U10AA008401).

